# In vitro effect of a non-immunosuppressive FKBP ligand, FK1706, on SARS-CoV-2 replication in combination with antivirals

**DOI:** 10.1101/2022.02.03.479080

**Authors:** William E. Fitzsimmons, Tracy L. Hartman, Michelle Mendenhall, Catherine Z. Chen

## Abstract

FKBP, a naturally occurring ubiquitous intracellular protein, has been proposed as a potential target for coronavirus replication. A non-immunosuppressive FKBP ligand, FK1706, was studied in vitro in a Vero cell model to assess potential activity alone and in combination with antivirals against SARS-CoV-2 replication. When combined with remdesivir, synergistic activity was seen (summary synergy score 24.7±9.56). FK1706 warrants in vivo testing as a potential new combination therapeutic for the treatment of COVID-19 infections.

---

FKBP is one of the naturally occurring ubiquitous intracellular proteins called immunophilins that has enzymatic activity as a peptidyl prolyl cis-trans isomerase and is also essential to the pharmacologic activity of immunosuppressants. The binding of tacrolimus, everolimus, and sirolimus, to FKBP is necessary but not sufficient to produce immunosuppression (1,2).

Replication of human coronaviruses is dependent on active immunophilin binding and inhibition of cyclophilins, an intracellular immunophilin, by cyclosporine blocks the replication of CoVs of all genera tested, including SARS-CoV, human CoV-229E and -NL-63, feline CoV, as well as avian infectious bronchitis virus (3-6). More recently, the immunophilin FKBP has been described as one of the potential targets for SARS-CoV-2 (7,8).

Two ligands to FKBP that are not immunosuppressive, FK1706 (9,10) and ElteN378 (11,12) were studied. These compounds are structurally distinct; both bind to the core structure for FKBP but do not have intact calcineurin or mTOR binding domains that produce immunosuppression. Because these drugs target host cells and may work by a unique mechanism to inhibit coronavirus replication, the additive or synergistic effect with known virus-targeting antivirals with mechanisms of RNA polymerase inhibition (e.g., remdesivir), viral error catastrophe or viral lethal mutagenesis (e.g., molnupiravir), or protease inhibition (e.g., M128533) were evaluated.

Vero E6 cells were infected with the live SARS-CoV-2 virus (USA-WA1/2020; World Reference Center for Emerging Viruses and Arboviruses (WRCEVA)) at low MOI (multiplicity of infection) and multiple rounds of viral replication occurred over the course of the assay. Percent CPE in compound-treated virus-infected cells were normalized to infected untreated cells as 0% and uninfected cells as 100% CPE protection. Based on these data, a concentration-response curve was created. Toxicity was assessed and compared in untreated, uninfected cells compared to treated cells.

In vitro testing was conducted at two independent laboratories in sequence. The details of the protocol followed by each laboratory are included in the appendix materials.

FK1706 (Shanghai SIMR Biotechnology Co. LQY20200910), ElteN378 (Glixx Laboratories Inc. GLXC -20448), remdesivir, molnupiravir, and M128533 were solubilized in DMSO and were diluted in culture test media to prepare compound concentrations.

Synergy was calculated using SynergyFinder 2.0 software (13). A summary synergy score greater than 10 was considered synergistic.

The initial results of FK1706 alone and in combination with remdesivir, molnupiravir, and M128533 are summarized in Table 1.

**Table 1.**
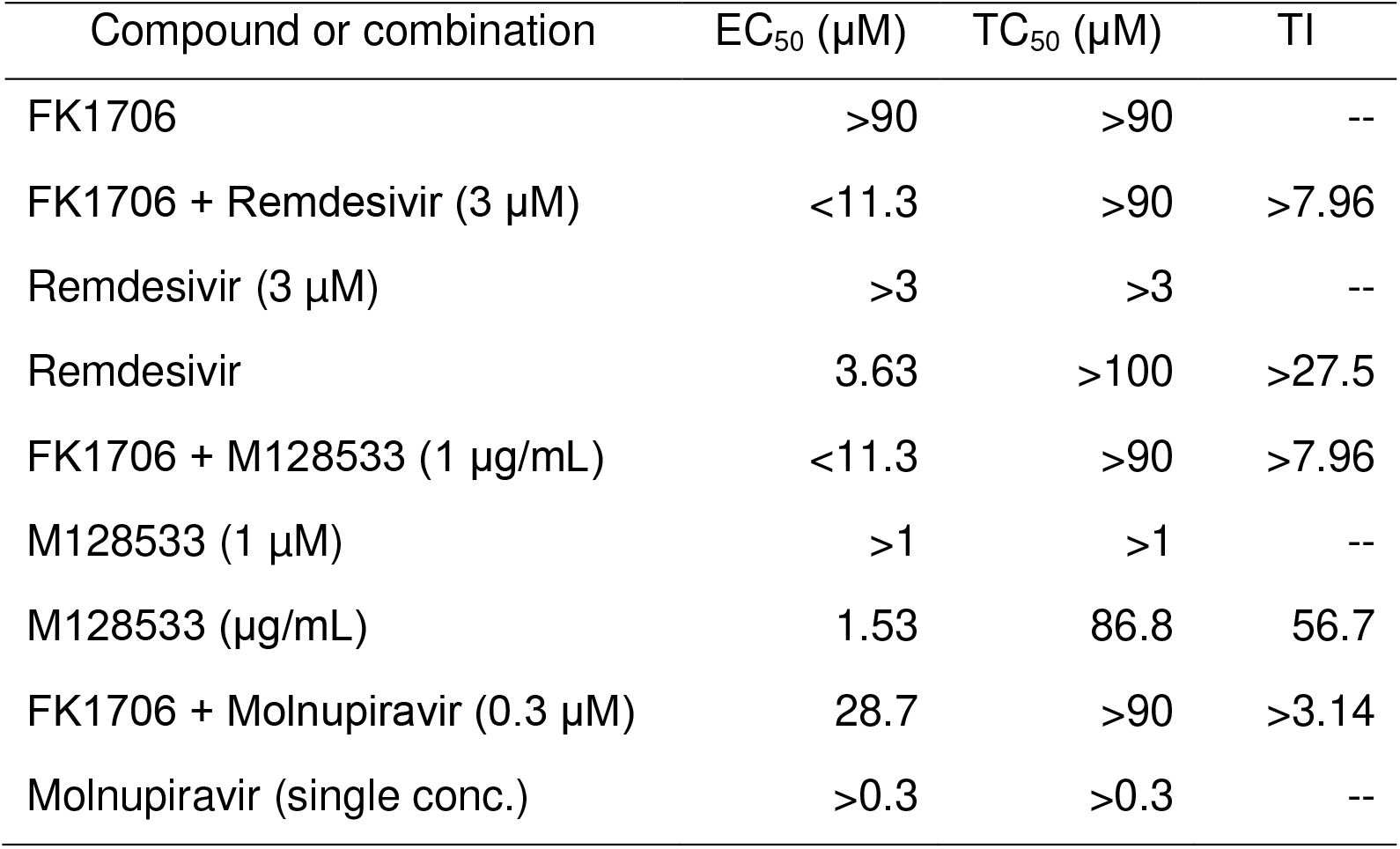
Anti-SARS-CoV-2 Cytoprotection Assay Results for FK1706 and antivirals against SARS-CoV-2 (USA-WA1/2020).

| Compound or combination | EC<sub>50</sub> (μM) | TC<sub>50</sub> (μM) | TI |
| --- | --- | --- | --- |
| FK1706 | >90 | >90 | -- |
| FK1706 + Remdesivir (3 μM) | <11.3 | >90 | >7.96 |
| Remdesivir (3 μM) | >3 | >3 | -- |
| Remdesivir | 3.63 | >100 | >27.5 |
| FK1706 + M128533 (1 μg/mL) | <11.3 | >90 | >7.96 |
| M128533 (1 μM) | >1 | >1 | -- |
| M128533 (μg/mL) | 1.53 | 86.8 | 56.7 |
| FK1706 + Molnupiravir (0.3 μM) | 28.7 | >90 | >3.14 |
| Molnupiravir (single conc.) | >0.3 | >0.3 | -- |

When combined, FK1706 (11-90 μM) and remdesivir (3 μM) were effective in inhibiting SARS CoV-2 viral CPE (93-100%, see Appendix Fig A1). FK1706 (2.85-90 μM) and molnupiravir (0.3 μM) inhibited SARS CoV-2 CPE (up to 70% reduction in viral CPE at 90 μM FK1706 with 0.3 μM molnupiravir; see Appendix Fig A2). FK1706 (11-90 μM) and M128533 (1 μg/mL) reduced SARS CoV-2 CPE (64-100%, see Appendix Fig A3).

Although FK1706 alone did not exhibit inhibitory activity against SARS-CoV-2, when combined with suboptimal concentrations (less than the EC_50_) of all three antivirals, increased inhibition was observed. Additive effects of ElteN378 with either remdesivir or M128533 were also demonstrated (see Appendix Table A1).

In follow-up confirmatory combination studies, FK1706 at multiple concentrations was tested in combination with multiple concentrations of remdesivir. When combined, remdesivir and FK1706 exhibited synergistic activity inhibiting SARS-CoV-2 and shifting the EC50 value of both compounds when in combination with the other (Figures 1A,B). The summary synergy scores were 24.7±9.56 by the ZIP (Supplementary Fig A4,A5), 24.8±9.56 by the Bliss and 24.9±9.56 by the HSA models. Scores >10 in all 3 models indicate synergy.

**Figure 1A.**
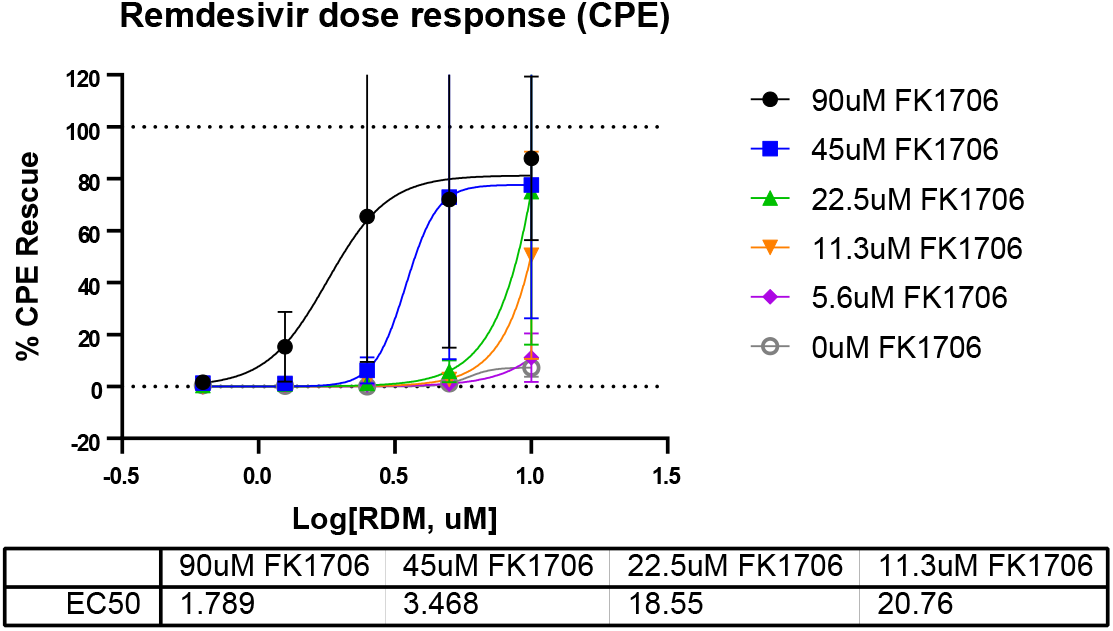
Concentration response of FK1706 when combined with remdesivir (RDM).

**Figure 1B.**
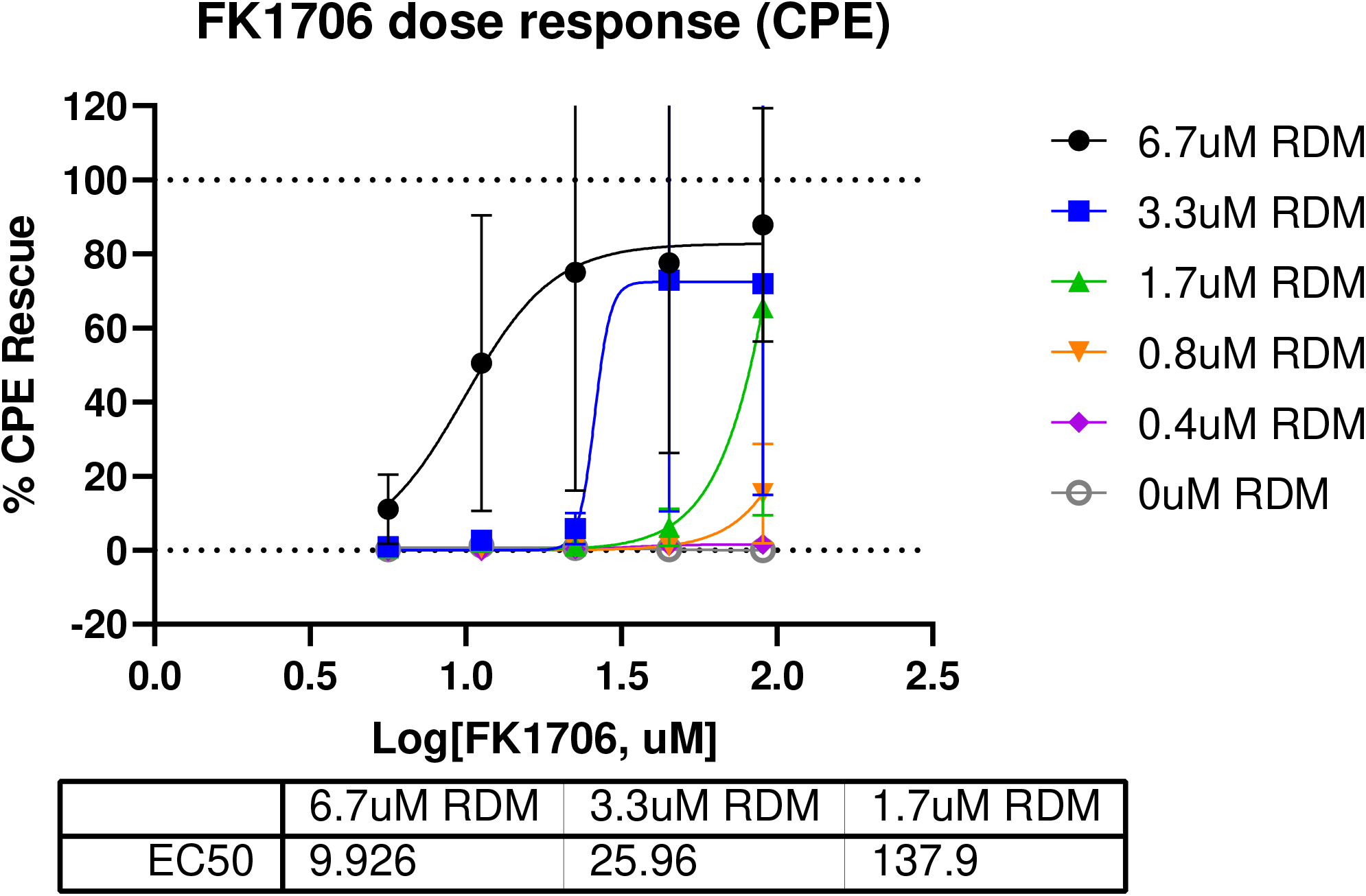
Concentration response of remdesivir when combined with FK1706.

Molnupiravir alone, nor in combination with FK1706, did not demonstrate activity in the follow-up confirmatory study. There was no evidence of cytotoxicity with FK1706 or remdesivir alone or in combination (Appendix Figs A6,A7).

The synergistic effects of FK1706 in combination remdesivir were demonstrated in a live SARS-CoV-2 virus assay measuring the ability of compounds to inhibit viral-induced CPE in Vero E6 host cells in vitro. The CPE reduction assay is a popular and widely used assay format to screen for antiviral agents because of its ease of use in quantitative high-throughput screening. The CPE reduction assay indirectly monitors the ability of compounds to inhibit viral replication and infection through various mechanisms, including direct inhibition of viral entry or enzymatic processes as well as acting on host pathways that modulate viral replication. This assay was previously used to screen 8,810 approved and investigational drugs from the National Center for Advancing Translational Sciences (NCATS) small molecule collections (14). A cytotoxicity counter-screen was conducted in parallel in host cells without addition of virus and demonstrated no substantial cytotoxicity of any of the test agents alone or in combination.

Since two chemically distinct FKBP ligands, FK1706 and ElteN378, both demonstrated activity, it is likely that FKBP is the key target. This target is in the host cells and complements the virus-targeted antivirals. The combination activity of these FKBP ligands was not limited to a single virus-targeted mechanism as the three antivirals have distinctly different mechanisms.

Remdesivir (Veklury), currently the only FDA-approved antiviral for COVID-19 infections, is administered intravenously to patients (15). Molnupiravir has received Emergency Use Authorization as oral therapy for outpatient COVID-19 infections (16). Although both of these antivirals have demonstrated clinical efficacy, there is a need for higher response rates and FK1706 may have utility in both settings. Additionally, these combinations should be active against variants with mutations in spike protein.

Both live virus assays use Vero E6 as host cells. Vero E6 cells have been shown to have high drug efflux transporter P-glycoprotein (P-gp) activity, which can reduce cellular concentrations of test articles, and remdesivir is a known P-gp substrate (17). Therefore, synergy observed in Vero E6 cells could be due to P-gp inhibition, which enhances the exposure of remdesivir *in vitro*, and warrants repeating in other cell-based models.

FK1706 has completed all nonclinical safety pharmacology, ADME, and GLP toxicity studies to support clinical development. Phase 1 healthy volunteer and Phase 2 studies in patients with neuropathy have been completed (9). This clinical experience would expedite the introduction of FK1706 into clinical studies of patients infected with SARS-CoV-2.

In conclusion, these data demonstrate that FKBP is a valid target for coronavirus infections in combination with virus-targeted antivirals such as remdesivir and molnupiravir. FK1706 warrants testing in an in vivo animal model of SARS-CoV-2 and if promising, rapid introduction into COVID-19 infection clinical trials.

## Acknowledgements

The authors thank Diane M. Coniglio, Pharm.D., President, Opus Medical Communications for editorial assistance with the manuscript.

