## Appendix Figures, Methods, Table for "In vitro effect of a non-immunosuppressive FKBP ligand, FK1706, on SARS-CoV-2 replication in combination with antivirals"

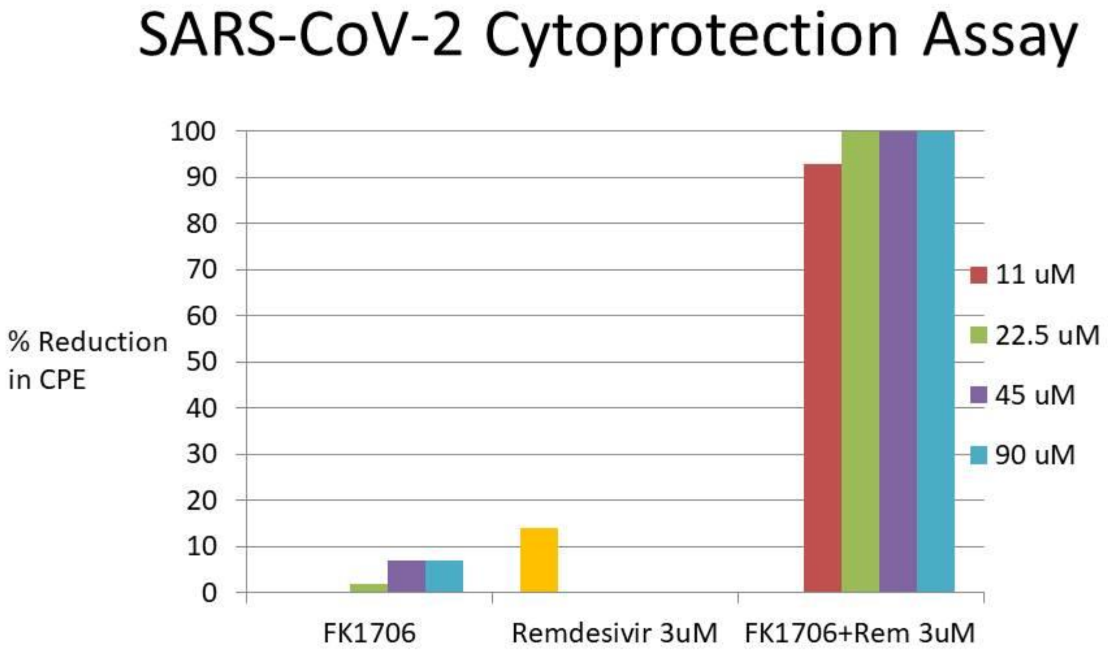


**Figure A1. FK1706 concentration response and remdesivir (Rem) at 3 uM**

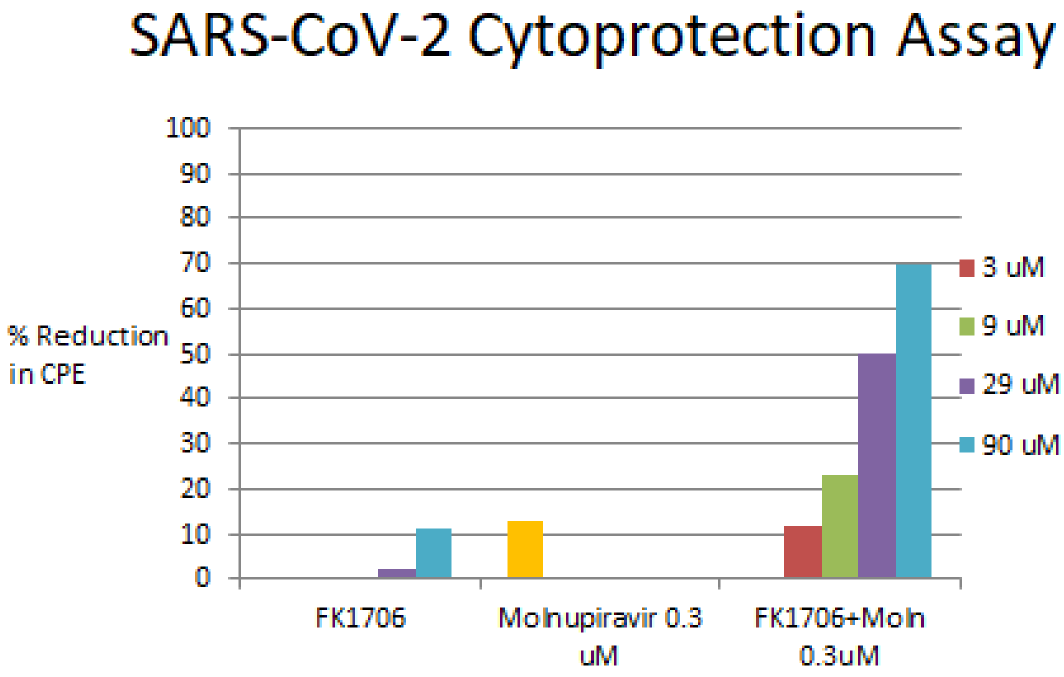


**Figure A2. FK1706 concentration response and molnupiravir (Moln) at 0.3 uM**

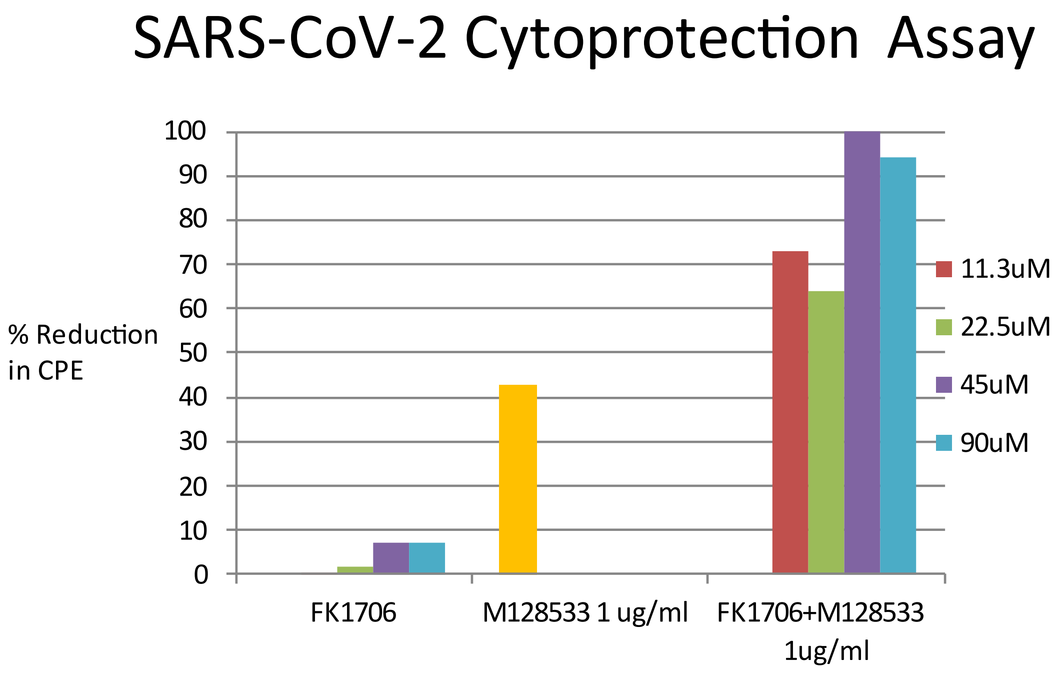


**Figure A3.** **FK1706 concentration response and M128533 at 1 ug/ml**

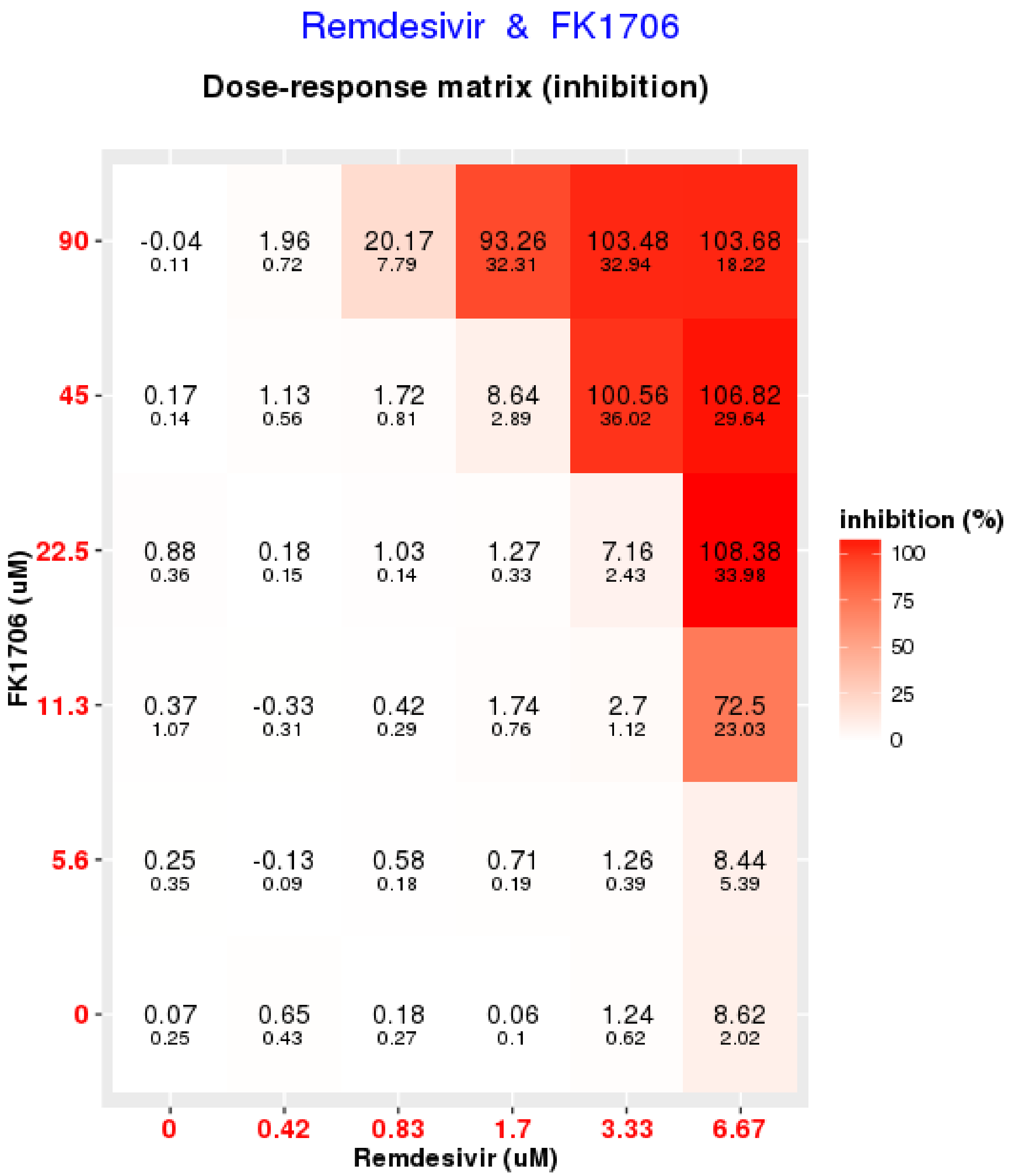


Figure A4. Concentration response of ZIP model synergy scores

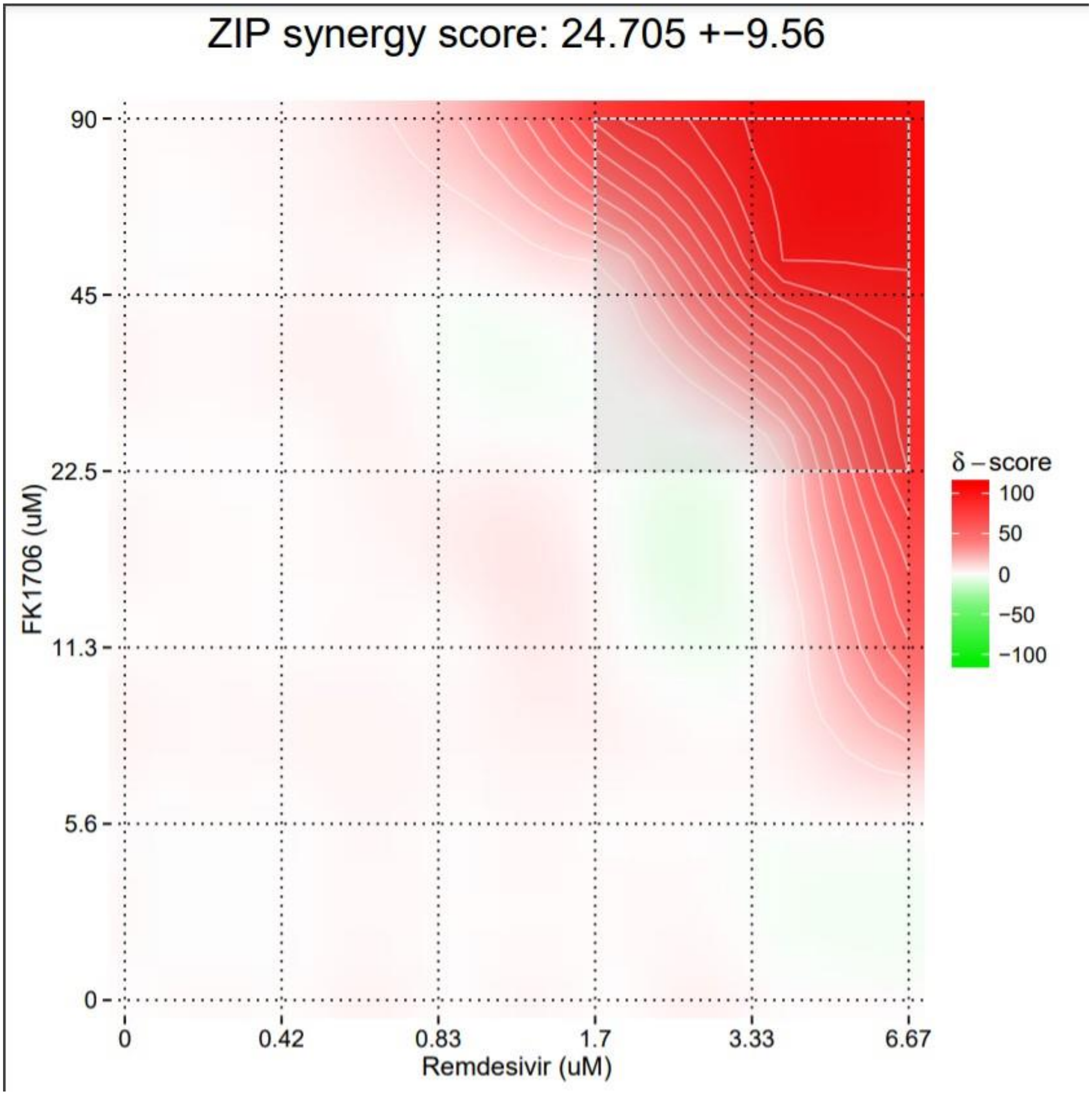


Figure A5 Synergy Score using the ZIP model

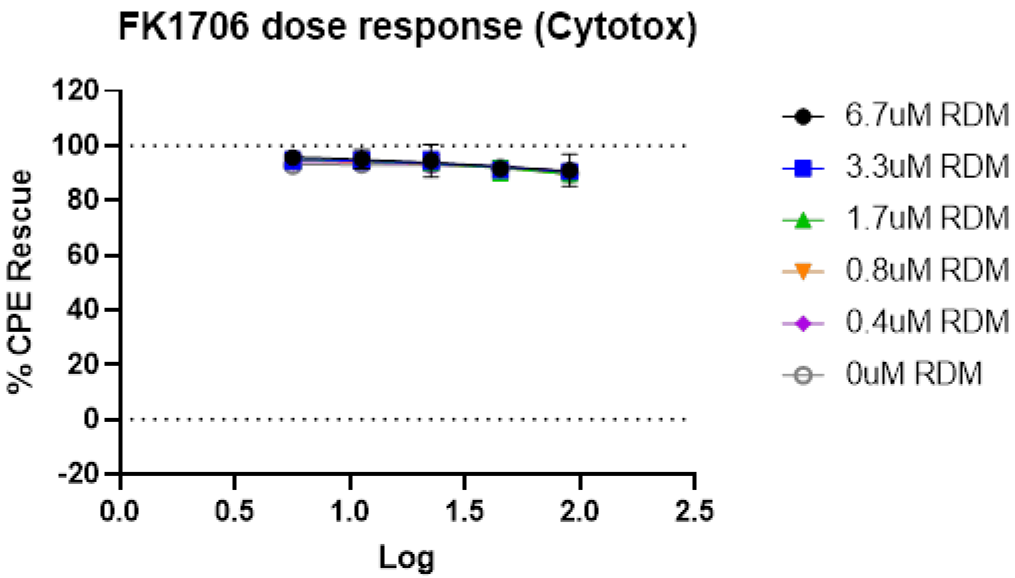


**Figure A6. FK1706 Cytotoxicity concentration response. RDM = remdesivir**

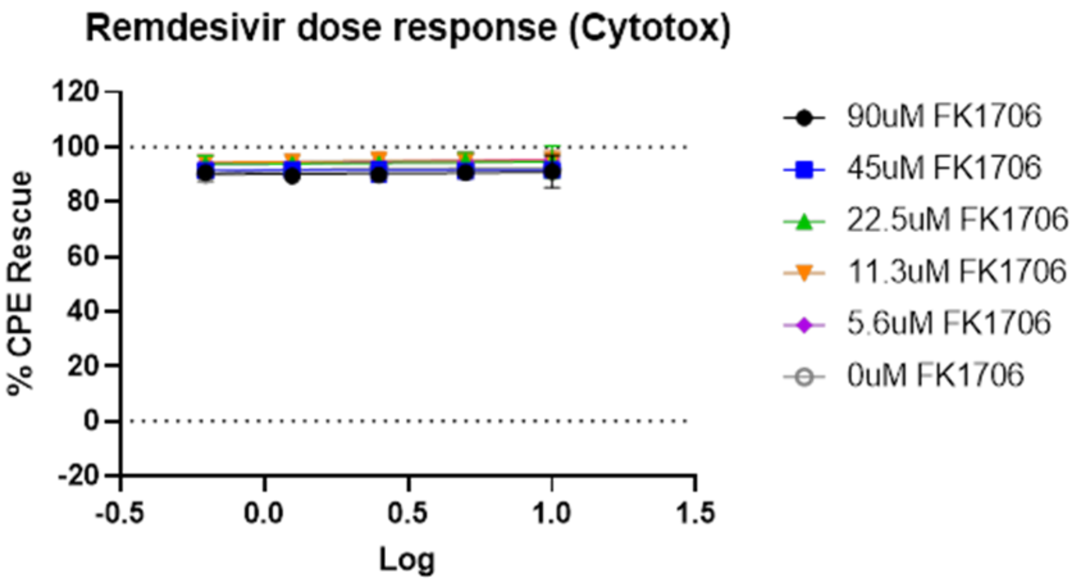


**Figure A7. Remdesivir cytotoxicity concentration response.**

### Methods

#### Initial Testing at the Antiviral Research Institute, Utah State University

The test compounds included FK1706, ElteN378, remdesivir, molnupiravir and M128533. These test compounds from DMSO stocks were serially diluted in MEM solution with 50 µg/mL gentamicin and 2% FBS. Each dilution was added to 5 wells of a 96-well plate with 80-100% confluent Vero E6 cells (3 x $10^4$ cells per well). SARS-CoV-2 (USA-WA1/2020) virus was prepared to achieve the lowest possible multiplicity of infection (MOI; approximately 0.001) that would yield >80% cytopathic effect (CPE) at 3 days. Three wells of each dilution were infected with virus and two wells remained uninfected as toxicity controls. Protease inhibitor M128533 was tested in parallel as a control compound. After untreated virus control wells reached maximum cytopathic effect (CPE) following 3 days' infection, plates were stained with neutral red dye for approximately 2 hours. Supernatant dye was removed and incorporated dye was extracted in 50:50 Sorensen citrate buffer/ethanol then read on a spectrophotometer at 540 nm.

#### Confirmatory Testing at Southern Research Institute (SRI International)

Remdesivir, molnupiravir, and FK1706 were dissolved and titrated in DMSO. To prepare assay-ready plates, compounds were acoustically dispensed as 45 nL per compound per well (90 nL/well total DMSO volume) in 6x6 matrix combinations into 384-well assay plates (Greiner, black clear bottom, tissue culture treated plates). Each compound was titrated at 5-point 1:2 titrations with the sixth point being DMSO only (to test single agent dose response). Each matrix block was tested as n=3. 5 µl/well of media was dispensed into assay plates (MEM, 1% Pen/Strep/GlutaMax, 1% HEPES, 2% HI FBS). Host cells were Vero E6 cells selected for high ACE2 expression (Severson et al, 2007). 25 µL/well of Vero E6 cells inoculated with SARS CoV-2 (USA_WA1/2020) at an MOI of 0.002 suspended in media were dispensed. The final cell density was 4000 cells/well. Assay plates were incubated for 72 hours at 37°C, 5% $CO_2$, 90%

humidity then viability was assayed by Vero E6 host cell ATP content. 30 µL/well of CellTiter-Glo (Promega, Cat # G7573) was dispensed. Plates were incubated for 10 minutes at room temperature. Luminescence signal was read on Perkin Elmer Envision or BMG CLARIOstar plate reader.

Cytotoxicity was assessed in a cell-based assay measuring host cell ATP content as a readout for cytotoxicity in Vero E6 cells selected for high ACE2 expression (Severson et al, 2007). This assay is used as a counter screen assay to the SARS-CoV-2 CPE Assay.

Raw assay signals were converted to percent of cell controls and normalized to the virus control.  The concentration of test compound required to inhibit CPE by 50% ($EC_{50}$) was calculated by regression analysis. The concentration of compound that would cause 50% cell death in the absence of virus was similarly calculated ($TC_{50}$). The selective index (TI) is the $TC_{50}$ divided by $EC_{50}$.

**Table A1. Anti-SARS-CoV2 Cytoprotection Assay Results of ElteN378 with antivirals**

| **Compound(s)** | **VeroE6/SARS CoV-2 (USA-WA1/2020)** | | |
|---|---|---|---|
| | **$EC_{50}$ (µM)** | **$TC_{50}$ (µM)** | **TI** |
| ElteN378 | >30 | >30 | -- |
| ElteN378 + Remdesivir (3 µM) | 19.6 | >30 | >1.53 |
| Remdesivir | 7.31 | >100 | >13.7 |
| ElteN378 + M128533 (1 µg/mL) | <1.11 | >30 | >27 |
| M128533 (µg/mL) | 1.36 | >100 | >73.5 |

ElteN378 showed little reduction in SARS CoV-2 virus (0-7%) across the range of 0.12-30 µM. Remdesivir alone at a subtherapeutic concentration (i.e. 3 µM) also showed very little reduction in SARS CoV-2 virus.  When combined ElteN378 (0.12-30 μM) and remdesivir (3 μM) reduced SARS CoV-2 virus (up to 70% reduction in viral CPE at 30 μM ElteN378 with 3 μM remdesivir).

M128533 alone at a suboptimal concentration (i.e. 1 μg/mL) showed reduction in SARS CoV-2 virus ranging from 41-76%.  When combined, ElteN378 (0.12-30 μM) and M128533 (1 μg/mL) further reduced SARS CoV-2 virus CPE (69-95%).

These data demonstrate that ElteN378 combined with subtherapeutic remdesivir or M128533 suppresses SARS-CoV-2 virus replication.
